# Motor Cortex Projections To Red Nucleus And Pons Have Distinct Functional Roles In The Mouse

**DOI:** 10.1101/2022.09.06.506669

**Authors:** Veronica Lopez-Virgen, Martin Macias, Paola Rodriguez-Moreno, Rafael Olivares-Moreno, Victor de Lafuente, Luis Concha, Gerardo Rojas-Piloni

## Abstract

Pyramidal tract neurons (PTNs) are fundamental elements for motor control. However, it is largely unknown if PTNs are segregated into different subtypes with distinct roles in movement performance. Using anatomical, electrophysiological and optogenetics tools, we analyzed in both sexes’ mice motor cortex, PTNs projecting to red and pontine midbrain nuclei, which are important hubs connecting cerebral cortex and cerebellum playing a critical role in the regulation of movement. We reveal that vast majority of M1 neurons projecting to the red and pontine nuclei constitutes different populations. Corticopontine neurons have higher conduction velocities and morphologically, a most homogeneous dendritic and spine distributions along cortical layers. Optogenetically inhibiting either kind projection, differentially affects forelimb movement onset and execution in a lever press task, but only the activity of corticopontine neurons is significantly correlated with trial-by-trial variations in reaction time. The results indicate that cortical neurons projecting to the red and pontine nuclei constitute distinct functional and anatomical pathways and they contribute differently to sensorimotor integration, suggesting that layer 5 output neurons are functionally compartmentalized generating, in parallel, different downstream coding.

## 1. INTRODUCTION

Pyramidal tract neurons (PTNs) of the sensorimotor cortex are essential for motor control transmitting information directly to several subcortical structures including the spinal cord (Olivares-Moreno et al., 2017), medulla (Oswald et al., 2013), thalamus (Jiang et al., 2020), hypothalamus (Winnubst et al., 2019) and mesencephalic nuclei (Economo et al., 2018). PTNs are thick-tufted cells located in layer 5b of sensorimotor cortex that differ from intratelencephalic slender-tufted pyramidal neurons located in layer 5a, which project mainly to the striatum and contralateral cortex (Oberlaender et al., 2011). Despite PTNs has been extensively studied, is largely unknown if they are segregated into different subtypes executing in parallel specific computations important for movement performance.

Although several studies have shown that PTNs project almost individually to their respective targets (Akintunde and Buxton, 1992; Groh et al., 2010; Oswald et al., 2013; Rojas-Piloni et al., 2017), others have found that PTNs do not segregate into different projection types (Kita and Kita, 2012; Guo et al., 2017) with several collaterals of a single cell synapsing into more than one target. For example, PTNs projecting to subthalamic nucleus also send collaterals to the striatum, associative thalamic nuclei, superior colliculus, zona incerta, pontine nuclei (PN), multiple other brainstem areas, and the spinal cord (Kita and Kita, 2012). Additionally, Guo et al. (2017) reported a high multidirectional projection index of L5 thick-tufted neurons, including corticospinal and corticostriatal neurons projecting to the midbrain, thalamus, and hypothalamus. This suggests that the sensorimotor cortex transmits parallel copies of information to several areas, but certain zones receive specific corticofugal information. In this way, recently (Economo et al., 2018) it has been found that corticothalamic and corticoreticular PTNs constitute distinct neuronal populations, with specific gene expression and electrophysiological activity related to tongue movement. However, neurons projecting to the midbrain, corticopontine and corticotectal nuclei seem to display fewer particularities (Rojas-Piloni et al., 2017; Economo et al., 2018).

Two important mesencephalic zones receive direct cortical inputs, the red nucleus (RN) and pontine nuclei (PN) and both play a fundamental role in motor control, forming loops of information between the cortex and cerebellum in rodents (Liang et al., 2012; Wagner et al., 2019) and primates (Lemon, 2016). The parvocellular part of the RN receives ipsilateral projections from cortical neurons of sensorimotor areas (Humphrey and Rietz, 1976; Catman-Berrevoets et al., 1979), which in turn project to the ipsilateral lower olive and then to the cerebellum (Massion, 1967; Edwards, 1972; Walberg and Nordby, 1981; Kennedy, 1990). The magnocellular RN receives information from the nucleus interpositus of the contralateral cerebellum (Jansen and Jansen Jr, 1955; Angaut and Bowsher, 1965; Courville, 1966; Daniel et al., 1988) and is the main origin of the rubrospinal tract (for a review, see (Olivares-Moreno et al., 2021). On the other hand, PN receive ipsilateral inputs from several cortical areas in a topographical way (Schwarz and Thier, 1995; Huang et al., 2013), and then the pontocerebellar tract reaches most parts of the cerebellum, constituting the major afferent input to the cerebellum (Brodal, 2014).

To understand the organization of PTNs and if different functional subtypes contribute with specific computations, here we analyzed functional and morphological aspects of two classes of PTNs projecting to the RN and PN. The results indicate that, even though are topographically intermingled, distinct types of PTNs differentially modulate the RN and PN, suggesting that layer 5 output neurons are functionally compartmentalized.

## MATERIAL AND METHODS

### Animals

All procedures were carried out in strict accordance with the recommendations of the National Institutes of Health Guide for the Care and Use of Experimental Animals and the Laboratory Animal Care (Mexican Official Norm [NOM] 062-ZOO-1999); the procedures were approved by the local Animal Research Committee of the Instituto de Neurobiología at Universidad Nacional Autonoma de Mexico (UNAM). We used eight-week-old male or female C57BL/6 mice that were maintained at constant room temperature (22 ± 2°C) under a 12-h light/dark cycle.

### Retrograde tracer injections

To quantify the number of corticorubral (CR) and corticopontine (CP) PTNs, we injected two retrograde neuronal tracers into the RN and PN. The animals were anesthetized with isoflurane/O_2_gas (1.5%) and placed in a stereotaxic frame (World Precision Instruments, Inc., cat #E04008-005) and given an injection of 2% lidocaine (0.10 cc, s.c.) at the incision site. Then, a 1.5 cm incision across the midline was performed to expose the skull and identify the bregma suture. Small craniotomies were done with a dental drill over the injection sites using the following coordinates: RN 3.4 mm posterior from bregma, 0.7 from midline and 4 mm depth; PN 4 mm posterior from bregma, 0.5 mm from midline and 5.5 mm depth from the pia. Retrograde tracers were pressure injected (80-100 nL) using a picopump (WPI Inc, PV830 Pneumatic PicoPump) coupled to a calibrated glass injection capillary (BLAUBRAND® intraMARK REF 7087 07). The volume delivered by capillary was supervised visually in each injection. After injection of tracers, the incision site was thoroughly cleaned with saline and sutured. Injections into RN and pons of the same animal were performed using a combination of the retrograde tracers Fluorogold (FG) (Fluorochrome, LLC; 3% in distilled water) and Dextran tetramethylrhodamine 3000 MW (BDA) (Molecular Probes; 1 mg/ml in PBS, cat # D3308). Five days after the injections the mice were perfused transcardially with 0.1 M phosphate buffer (PB) followed by 4% paraformaldehyde in 0.1 M PB. The brains were extracted and post-fixed overnight in 50 ml of paraformaldehyde. Then the brain was cut coronally in the sensorimotor cortex at 50 μm in an automated vibrating Microtome (Leica VT1200S) and from the injection site. Slices were mounted with SlowFade Gold (Molecular Probes, cat. num. S36937) after. The size of the injection sites was estimated automatically using ImageJ software (V 1.50i) outlining the periphery of the zone stained with the tracer in the center of each injection and computing the transversal area. Only the experiments in which the injections were located within the RN and PN were used for neuronal quantification and in which the injection sizes were equal. For NeuN staining, brain sections were rinsed three times in 0.1 M PB and incubated in a blocking solution for 40 min. Then, the sections were incubated in the primary antibody mouse anti-NeuN (dilution 1:500; Millipore, Cat # MAB377) in a blocking solution at 4°C overnight. Sections were rinsed 3x with 0.1 M PB and incubated in 0.1 M PB containing 10% goat serum and conjugated secondary antibody (Alexa Fluor 647 anti-mouse, dilution 1:500, Invitrogen cat #A-21235) for 1 h at room temperature.

Mosaic images (resolution: 1.023 μm/pixel) of the sections containing the fluorescent retrograde-labeled cells were obtained on a fluorescence microscope (Zeiss AXIO Imager.Z1) attached to a digital camera (AxioCam MRm, 1.3 MP) using the appropriate filters (GFP for Alexa 488, Rhodamine for Alexa 594 and 650 nm for Alexa 647) and acquired with a 10x objective (ZEISS Plan-APOCHROMAT, NA: 0.45). Additional detailed images (resolution: 0.66 μm/pixel) were acquired with a confocal microscope (Zeiss 780 LSM) using an objective LD PCI Plan-Apochromat 25x/0.8 Imm Korr DIC M27.

### Analysis of the retrograde tracers

For the injection site quantification, we used ImageJ1.51u, and to analyze the projection neurons in the sensorimotor cortex. The microscopy images (1.023 μm/pixel) were aligned manually in Amira (version 5.6) coupled to magnetic resonance image volume (http://imaging.org.au/AMBMC/Model) by selecting shared and clearly visible anatomical landmarks and using a linear transformation (Janke & Ullmann, 2015). Then when the microscopy images were aligned, the positions of the PTN somas were labeled. In this way, a 3D map of the CR and CP neurons was obtained. To compute relative neuron density, the soma distributions were obtained in 100×100 mm steps for the coronal plane, and vertical density profiles were computed in 50 μm steps along the vertical axes in M1.

### In vivo electrophysiological recordings and stimulation

For electrophysiological experiments 71 mice were used. Mice were anesthetized with isoflurane/O_2_ gas mixture (1.5%), placed in a stereotaxic frame, and maintained at a constant temperature (37°C). A small craniotomy was made in the coordinates for RN (3.4 mm posterior from bregma, 0.7 from midline and 4 mm depth; n=34) or PN (4 mm posterior from bregma, 0.5 mm from midline and 5.5 mm depth from the pia; n=37), and a bipolar concentric electrode (MicroProbes CEA 200) was positioned using a micromanipulator” (Narishige SM-15L). Additionally, for the cortical recording of PTNs, a cranial window was made to expose M1 (0.98 mm posterior and 1 mm lateral relative to bregma). The *in vivo* juxtacellular recordings and biocytin fillings have been previously described in detail (Pinault, 1996; Narayanan et al., 2014). Briefly, recordings were performed with borosilicate glass pipettes (7-15 MΩ) and filled with normal rat Ringer’s solution with 2% biocytin (Sigma, cat # 576-19-2). The pipette was positioned with a micromanipulator (Luigs & Neumann, Mini Compact Unit). Electrophysiological signals were recorded in current-clamp mode using a microelectrode amplifier (Axon Instruments, Axoclamp 2B). Then, the output signal was amplified (x100) and filtered at a bandwidth of 30 Hz - 10 kHz with a differential amplifier (AM systems, model 1700). The resultant electrophysiological signals were digitized with a rate of 20 kHz using Digidata interface (1440A) and the pClamp software module, Clampex (V 10.3).

The pipette was advanced in 1 μm steps to locate single neurons, indicated by an increase in the electrode resistance (25-35 MΩ) and in the amplitude of action potentials (3-5 mV). Ongoing spiking was recorded for each neuron during a minimum of 60 s. For each recorded cell we tested if antidromic spikes were evoked following RN or PN stimulation. Antidromic spikes were tested using a single, 0.1 ms square pulse with intensities starting at 100 μA and increasing until a threshold was reached for each cell, but never exceeding 300 μA. When a stable cell recording was obtained, the following criteria were used to establish the antidromic characteristic of the cell responses: a constant threshold and latency, the ability to follow a stimulus train of 333 Hz and a collision of the orthodromic spikes with antidromic evoked spikes (Olivares-Moreno et al., 2017). We used the spontaneous action potentials of sensorimotor cortex recorded neurons to trigger the electrical stimulation with a variable delay. Systematically changing the delay between the spontaneous spikes and stimulation allowed us to measure the critical period in which collision between spontaneous and evoked action potentials occurs. Neurons that did not satisfy these criteria were considered non-identified neurons.

Following the recording, juxtasomal biocytin filling was performed by applying continuous, low intensity square pulses of positive current (<7 nA, 200 ms on/200 ms off, Master-8, A.M.P.I.), while gradually increasing the current in steps of 0.1 nA and monitoring the AP waveform and frequency. The membrane opening was indicated by a sudden increase in AP frequency. Filling sessions were repeated several times (15–30 min) and diffusion was allowed for 1–2 h to obtain high-quality fillings.

After the experiments, the tip location of the stimulating electrodes was assessed with electrolytic lesions (100 mA, 10 s). At the end of the experiments, the animals were perfused, and coronal sections of interest regions were cut as indicated above. Slices with electrolytic lesions were mounted and stained with Nissl for electrode location analysis.

### Histology and image acquisition

Animals were perfused transcardially with 25 ml of 0.1 M PB solution followed by 25 ml of 4% paraformaldehyde in 0.1 M PB solution at pH 7.4. The brain was extracted and post-fixed overnight in paraformaldehyde. In experiments with *in vivo* recordings and biocytin filling, the cortex was cut into consecutive 50 μm thick tangential slices and treated with Streptavidin Alexa-488 conjugate (5 mg/ml Molecular Probes, cat num S11223) in PB with 0.3% TX for 3–4 h at room temperature to stain biocytin-labeled morphologies. All slices were mounted on glass slides, embedded with SlowFade Gold (Molecular probes, cat. num. S36937) and enclosed with a cover slip.

### Image acquisition

For the 3D reconstruction of individual dendritic morphologies (biocytin-488 nm), the images were acquired in a confocal system (Zeiss 780 LSM) with a x63 oil immersion objective (Zeiss, EC Plan-Neofluar 40x/1.30 Oil DIC M27) with a 405 excitation laser (emission detection range: 400-455 nm). Mosaic images were acquired at a resolution of 0.109 μm x 0.109 μm x 0.5 μm per voxel for ∼30 consecutive 50 μm thick brain slices to cover complete dendrite morphologies from the pia to L6.

### 3D PTN reconstruction and quantitative morphology

For the 3D tracing and reconstruction of the dendrite morphologies of PTNs (without knowledge of the PTNs’ subcortical targets), image stacks were deconvolved using ImageJ software for confocal images (ImageJ 1.50i, USA). Then, the FilamentEditor of Amira software (Amira V 5.6) with the semi-automated method of (Oberlaender et al., 2007) were used and adapted for confocal microscopy (Rojas-Piloni et al., 2017). For the detection and automated reconstruction of dendritic spines, Neurolucida 360 software (Module AutoSpine 2.5 64-bits, Williston, Vermont, USA) was used.

Specific morphological characteristics of each reconstructed neuron were obtained (Table 2) and compared statistically for each group. The distribution of dendrite densities and dendritic spines every 50 mm from pia to L6 was statistically compared using the Kolmogorov-Smirnov test. Additionally, similarity between these distributions was computed as follows: 1) each distribution was normalized with respect to the respective maximum value; 2) the normalized distribution values of each individual PTN were subtracted from each of the two target-related average distributions. If subtraction resulted in negative values, the respective bins were set to zero; 3) for each resultant distribution, the bin-wise absolute difference between its normalized distribution and each of the two target-related distribution was calculated; 4) The bin-wise differences were summed across the entire depth (0-1000 μm). This sum was defined as the similarity between the dendrite or spines (i.e., the smaller the similarity value, the more similar the distributions). The similarity values obtained from dendrites or spine density distributions for each PTN class were combined to two similarity indices, as shown in Figure 3.

### Virus injection surgery

The activity of CR or CP neurons in M1 was analyzed with photometry during the motor execution task. To do so, we infected the neurons with the retrograde virus pGP-AAVrg-syn-jGCaMP7s-WPRE (Addgene, Plasmid #104487-AAVrg). Injections (450 μL) were performed similarly to the retrograde tracer injections (see above), but in these animals, an optical fiber cannula (Doric lenses, cat # MCF_200/250-0.66_1mm_ZF1.25(G)_FLT) was stereotaxically implanted in the left motor cortex (1.7 mm posterior and 1.5 mm lateral from bregma) below 500-600 μm from the pia and secured to the skull with dental cement (C&B Metabond®). For optogenetics, the retrograde virus AAVrg-hsyn-Jaws-KGC-GFP-ER2 (Addgene Plasmid # 65014-AAVrg) was injected into the RN or PN, and the optical fiber cannulas were implanted in the ipsilateral M1 cortex (600-1000 μm depth).

### Behavioral task

Ten days before the training, the mice were deprived of water (1 ml per day). The first training day the mice were placed in an operant conditioning box (18×16×15 cm) in which the lever was located at 1 cm from the floor, the light at 7 cm and the water dispenser at 2.5 cm. The lever was located to the right to force the animals to press with the right forelimb. The elements of the operant conditioned box were controlled with a custom-made platform (https://github.com/Juriquilla-ENES-INB-A13/SkinnerDuino-shield). In the initial phase, the mice were trained to press the lever and given a drop of water (8 μl) immediately after they did so. After that, the animals were trained to press the lever in response to a light. For this, the animals had to press the lever for the entire time the light remained ON (the maximum duration was 2 s). Immediately after the lever pressing, the light switched OFF, a drop of the reinforcer was delivered to the animal and a new trial started. The time interval between the trials varied randomly between 4 and 6 s. If the animals pressed the lever without the light, a 7 s time out would begin.

The animals learn the task in ∼10 sessions, reaching a plateau between 7 ± 0.6 of correct responses. The analysis of CR and CP neuron activity was performed in the sessions in which the animals reached their maximal efficiency.

### Photometry

The fluorescence emitted by CR or CP PTNs was detected with a photometry system (Multi-Fiber Photometry System, PLEXON Inc.). The system uses a blue light (465 nm LED) reflected by a dichroic mirror to excite GCamP7s. The emitted GCaMP7s fluorescence was recorded (Excitation wavelength 470 nm, Emission wavelength 505-545 nm) with CineLyzer software (Version 4.3.0 PLEXON Inc.). The trajectory of the lever pressings was recorded with a video camera (Integrated Imaging Solutions Inc. Model: FMVU-03MTC-CS) at 30 frames/s. Each trial lasted 5 s and started 400 ms before the cue (light) started. Forepaw movements for each trial (lever pressings) were analyzed offline with CineLyzer software (Version 4.3.0) to obtain the displacement during the time.

The bulk fluorescence emitted by CR and CP neurons in M1 and the simultaneous contralateral forelimb displacements were analyzed during 6 sessions per animal. Custom routines written in MATLAB were used for the analysis. Fluorescence is expressed as ΔF/F where F is the fluorescence intensity at any frame and ΔF is the difference between F and the resting fluorescence (minimum of the temporal average of raw fluorescence signal in a 3s window). The fluorescence and simultaneous lever displacement data were aligned to the beginning of the light and the beginning of movement (when the lever reached 3 mm), and then all trials per session were averaged. The latency between the beginning of the cue (light ON) and the peak of calcium activity was computed in each session. Additionally, the Pearson correlation coefficients between fluorescence peak time and the beginning of movement time for all the trials were computed.

### Optogenetics

Optogenetics photostimulation was performed with Radiant V2 2.2.0.10 PLEXON Inc.. Inactivation of CR or CP neurons during lever pressings was performed using pulses of orange light (620 nm; 90 mW/mm2) (Chuong et al., 2014) though optical fiber cannulas. Photoinhibition was applied in two types of trials (Figure 6B): 1) Before movement: ON starting with the light cue - OFF when animals press the lever (2 s in incorrect trials); 2) During movement: ON 800 ms after light cue starts with a duration of 2 s. The experimental sessions (6 per animal) consisted of 50% trials without and 50% photostimulation trials that were randomly distributed along each session. To compare the inhibition effectiveness, in 4 animals, we analyzed the effects in absence of the actuator using a retrograde virus that doesn’t express the halorhodopsin (AAVrg-hSyn-EGFP).

### Virtual tractography

To estimate the axonal length between M1 and either RN or PN, we queried the length of virtual tracts from the Allen Brain Connectivity Atlas (Oh et al., 2014). We selected 16 different axonal tracing experiments from the database with injection sites at M1 and targeting RN and PN (see list of tract tracing experiments below). The reconstructed streamlines were downloaded as json files through the streamline downloader service (https://neuroinformatics.nl/HBP/allen-connectivity-viewer/streamline-downloader.html) and converted to tck format using in-house routines built in Matlab R2018a. We combined all the streamlines from the 16 experiments into a single tractography file. Next, we used MRtrix3 (Tournier et al., 2019), a suite of tools built for diffusion-weighted MRI tractography to virtually dissect streamlines reaching either the ipsilateral RN or PN. The Allen Mouse Brain Atlas (Wang et al., 2020) was used to create binary masks of M1, RN and PN on the side of the injection site (Figure 3A). The resulting masks were used as inclusion criteria to dissect the tractography from M1, with three different criteria: [1] Streamlines reaching the ipsilateral PN (streamlines were truncated if they extended beyond PN, as they typically reached and arborized at the cerebellum); [2] streamlines reaching the RN by descending through the internal capsule then arching upwards to reach the ventral aspect of RN; and [3] streamlines reaching RN traversing the thalamus. Finally, we estimated the length of these pathways as the average distance of the streamlines included in each bundle.

Tract tracing experiments from the Allen Mouse Brain Connectivity Atlas.

180720175, 127084296, 100141780, 288169135, 166082842, 584903636, 126909424, 100141273, 277957908, 512130198, 298273313, 606250170, 272697944, 297711339, 156786234, 298325807, 100141563, 597007858, 168229113, 179641666, 166461193, 179640955, 182616478, 287461719, 177893658, 159651060, 310194040, 591612976, 157909001, 477836675.

### Experimental design and statistical analyses

Statistical analyses were computed using nonparametric tests. For multiple comparisons, a Kruskal-Wallis ANOVA and Friedman test were performed. Differences were considered significant starting at p=0.05. Calcium activity of PTNs was analyzed comparing ROC (receiver operating characteristic) curve analysis for trial-to-trial calcium signals comparing distributions of the fluorescence intensity each time with basal calcium activity (Tan, 2009). To analyze the projection neuron profile in the motor cortex, and for the physiological data, we used the Shapiro-Wilk test to determine in every case if the data were normally distributed, and we applied the appropriate statistical test to establish statistical differences (one-way ANOVA or Kruskal-Wallis and Bonferroni or Tukey post hoc tests, respectively) with Sigma Plot v12. To analyze the length and spines across the cortical depth we used a Kolmogorov-Smirnov test. Fluorescence values were compared with the control. Data are expressed as median and interquartile range, or the average and standard error. The level of confidence was set at 95% (p < 0.05).

## RESULTS

### Activity of cortico-rubral and cortico-pontine neurons during the execution of forelimb movements

As a first approximation, we use photometry to analyze if CR and CP PTNs are modulated in a specific manner during movement execution. Thus, we injected a retrograde viral vector (pGP-AAVrg-syn-jGCaMP7s-WPRE) into the RN or PN to enable the GCamp7s calcium indicator expression specifically in CR or CP PTNs. Additionally, an optical fiber cannula was implanted into M1 to measure by photometry the bulk calcium activity of subcortical projection neurons. No differences were found between the number of CP (238.01±22.7 neurons/ mm^2^) and CR (216.2±16.5 neurons/ mm^2^) expressing GCamp7s (t=-0.768; p=0.455). The animals were trained to press a lever in response to a light (Figure 1D) to receive a reinforcer (water). The animals (n=6) learned the task in approximately 11 sessions, reaching an efficacy (proportion of correct responses) of 68.2± 4.8% (Figure 1E). Once the animal reached stable behavior, we measured the changes in fluorescence in each correct trial. Changes in fluorescence were aligned to the beginning of a visual cue (lights ON) and to movement initiation. A significant correlation between calcium peak time and trial-by-trial lever press time was found for CP neurons but (r=0.22, p < 0.01) not for CR neurons (r=0.017, p=0.75), indicating that pons-projecting neurons synchronize their activity more precisely during the task.

**Figure 1.**
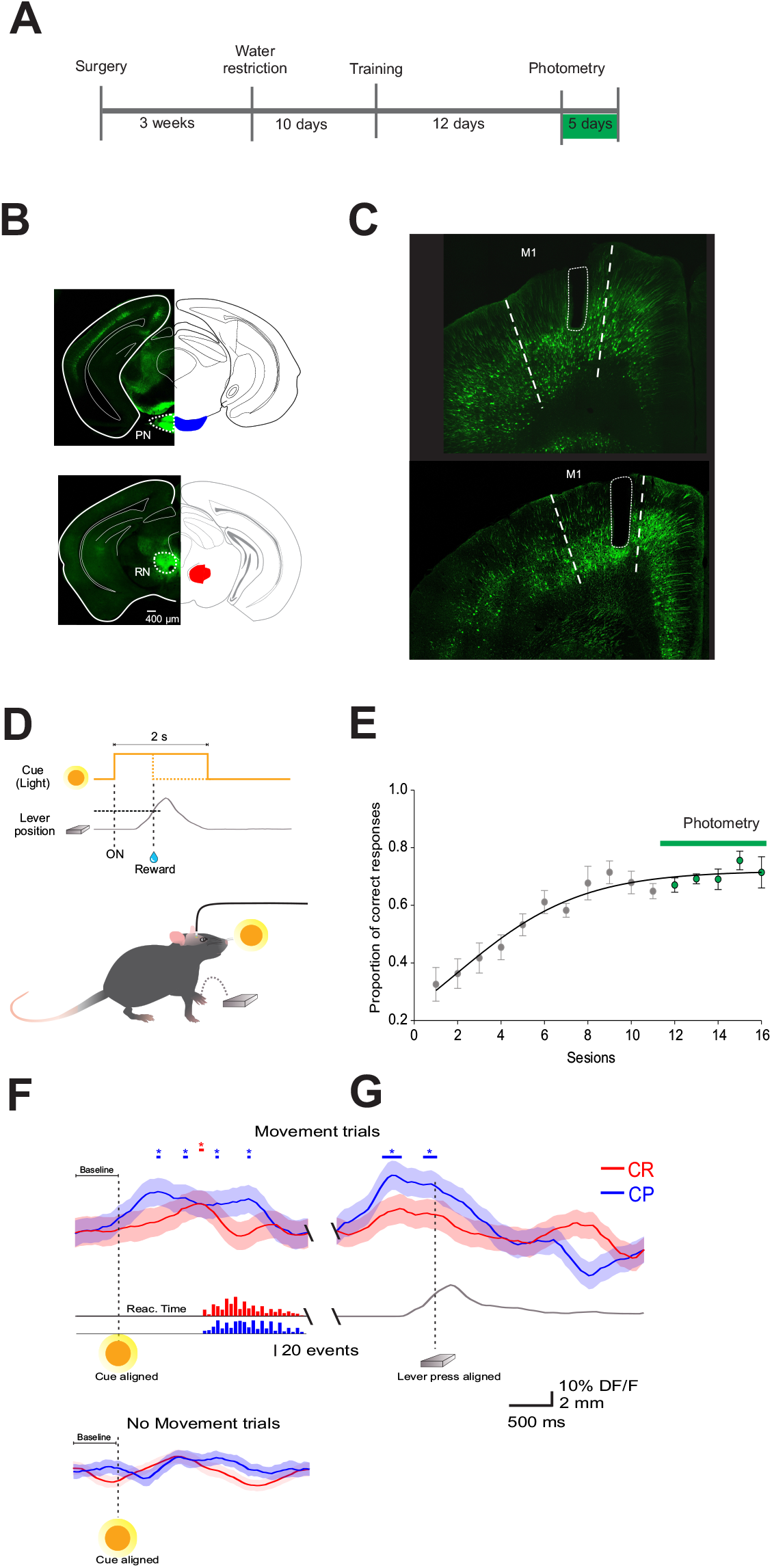
CR and CP PTNs are differentially modulated during execution of forelimb movements. **A**, Experiment timeline indicating the procedures. Surgery includes retrograde virus injection into the RN and PN, producing the expression of GCaMP7s in CR or CP PTNs, and optical fiber implantation. **B**, Exemplary viral injection sites into the PN (upper) and RN (lower). **C**, Coronal microphotography showing the expression of the retrograde virus in CP (upper) and CR (lower) neurons, as well as the optical fiber location in M1 (dashed area). **D**, Behavioral task: the animals learned to press the lever in response to a cue (light). The cue remained ON for 2 s, and the animals had to press the lever during this time to receive a reinforcer (drop of water). When the lever reached a threshold (dashed gray line; 3 mm) to deliver the reinforcer, the light was immediately turned OFF. The position of the lever was tracked with a video camera and digitized offline (gray trace). **E**, The animals learned the task in 10 days, reaching an efficacy (correct responses/total trials per session) of nearly 70%. In the last 5 sessions, calcium activity photometry for CR and CP neurons was recorded. **F**, Grand average (mean: lines; SE: shaded areas) of calcium fluorescence changes of CR (red) and CP (blue) aligned to the cue (vertical dashed line). Horizontal red (CR) and blue (CP) lines and asterisks above the traces indicate the time intervals in which fluorescence values increases significantly compared with baseline. (*p<0.05 area under the ROC curve). The histograms bellow shows the distribution of the reaction times computed for all trials. **G**, the same as F, but the fluorescence traces were aligned to the movement (threshold reached by the lever). The lower trace is the averaged displacement of the lever computed for all the trials in the photometry sessions. Traces bellow show calcium activity during omissions (trials without lever pressing).

To analyze the precise times in which calcium signal increases significantly for each type of neuron, we analyzed ROC (receiver operating characteristic) curves for trial-to-trial calcium signals to compare the distribution of fluorescence values for all the trials of basal activity (before cue). Basal fluorescence has been compared with distributions of the fluorescence intensity values from calcium signals each time. Figure 1 shows the precise times for which each calcium signals increase significantly compared to its own basal activity (horizontal lines upper the traces). The analysis reveals a different temporal dynamic between CP and CR neurons; only CP neurons significantly increased their activity before lever pressing movement (−533 to +33 ms respect to movement initiation). No significant increments of calcium signals were observed during omissions (trials without lever pressings).

### Anatomical distribution of cortico-rubral and cortico-pontine neurons

The previous results suggest that both classes of PTNs integrate different information and are compatible with the idea that PTNs that project to RN or PN might have different roles in the control of movement. Then, we analyzed the detailed distribution of CR and CP neurons in sensorimotor cortex. Two retrograde neuronal tracers were injected into the RN and PN (FluoroGold and BDA conjugated with rhodamine, respectively). Five days after injection, the animals were perfused for histological processing of the brain. The distribution of labeled cells was analyzed in 3 animals that were injected at both sites (Figure 2A, RN = 0.78 ± 0.081 mm^2^, PN = 0.82 ± 0.03 mm^2^, U = 97.00; p = 0.374, Mann-Whitney U statatistic). Additionally, to obtain a 3D representation of the PTN density, the histological mosaic images were aligned with the MRI images of the mouse brain atlas (http://imaging.org.au/AMBMC/Model (Ullmann et al., 2015). The results showed that both types of neurons are evenly distributed and intermingled in the rostro-caudal axis comprising the areas M1, M2, S1 and S2 (Figure 2D, F-G). but the vast majority of labeled neurons are those corresponding to the cortico-pontine projection (∼80%; M1 F = 8.096, p = 0.010; M2 F = 11.173, p = 0.004; S1F =19.628, p = <0.001; S2 F = 18.088, p =<0.001, One way ANOVA, Table 1). Interestingly, the proportion of neurons projecting simultaneously to the RN and PN is less than 4% in all cortical areas (Table 1). To validate if both neuronal retrograde tracers can be used simultaneously for the analysis of the relative neuronal densities of projection neurons, we inject both tracers (FG and BDA) into the same area of the spinal cord. In this way, we observe a large proportion (81.3±16.8, n=3 experiments) of cells with BDA also are positive for FG, indicating that the existence of double-labeled cells could be due to neuronal co-termination.

**Table 1.** Percentage of CR and CP neurons in sensorimotor cortices.

|  | M1 | M2 | S1 | S2 |
| --- | --- | --- | --- | --- |
| **Corticorubral** | 18.9 ± 6.9 | 17.4 ± 6.7 | 15.5 ± 6.7 | 15.8 ± 7.3 |
| **Corticopontine** | 78.1 ± 8.1 | 78.4 ± 8.3 | 81.4 ± 8.3 | 81.4 ± 8.8 |
| **Double labeled** | 2.9 ± 1.6 | 4.1 ± 1.7 | 3 ± 1.5 | 2.7 ± 1.6 |

**Figure 2.**
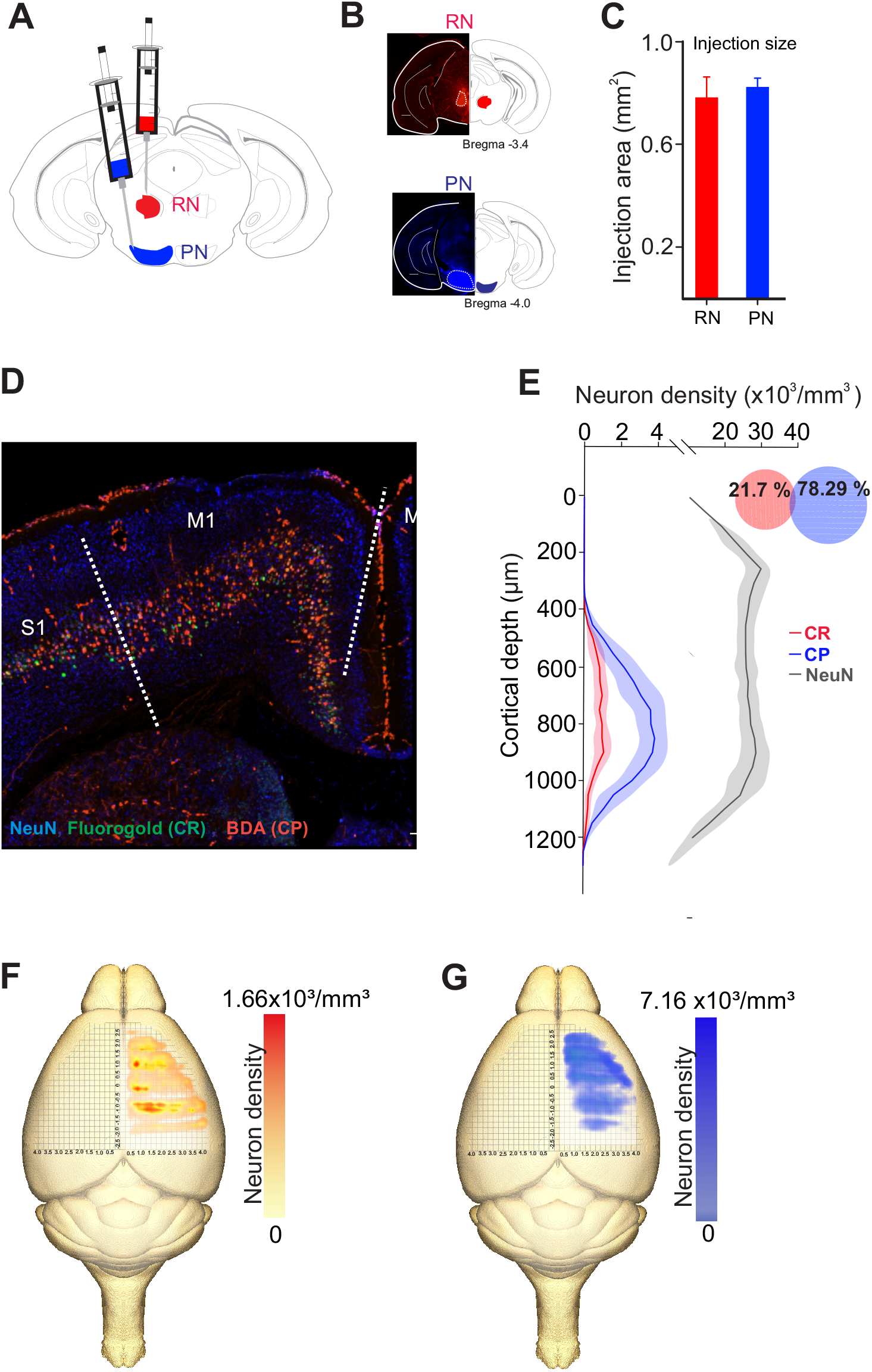
Distinct population of PTNs projecting to the RN and PN. **A**, Retrograde tracers FG and ChT 594 were simultaneously injected into the RN and PN, respectively. **B**, Examples of injection sites from the same animal. **C**, Comparison of the injection extent, measured as the transversal area covered by the tracer (dashed lines in B), in 3 mice (RN = 0.7 ± 0.08mm^2^, PN = 0.8 ± 0.03mm^2^). **D**, Coronal mosaic photomicrograph showing the distribution of CR (green) and CP (red) PTNs and NeuN (blue) in M1. **E**, Average vertical distributions of the CR (red) and CP (blue) PTN somata computed for 3 mice (3 consecutive slices per animal). Gray line represents the distribution of NeuN-labeled cells in the same experiments. Proportion of CR, CP and double-labeled neurons (0.7 %) in M1 are indicated in the Venn diagram. **F-G**, Averaged relative density representation maps (n=3 animals) in the horizontal plane of the CR (F) and CP (G) neurons. Neuron density is expressed as the mean number of retrograde labeled neurons (neurons/voxel 250 mm^3^).

Previous results have found that some PTN showed differences in their depth distribution along layer 5 (Groh et al., 2010; Rojas-Piloni et al., 2017; Economo et al., 2018) however, here we found that specifically in M1, CR and CP neurons have essentially the same cortical depth distributions (Figure 2E) and there in not a significant difference in the depth below the pia in both groups (CP: 819.1 ± 8.517 μm, n=393; CR: 789.4 ± 17.83 μm, n=113; p=0.1099, t=1.602, Two-tailed student t test).

### In vivo electrophysiological characteristics of cortico-rubral and cortico-pontine neurons

We next wondered whether neurons projecting to the RN and PN display distinct electrophysiological characteristics selective for their respective targets. For this, we performed single unit recordings of CR and CP neurons in M1, identified by the collision test of antidromic stimulation in the RN and PN (Figure 3A-B). Of the 242 neurons recorded, 45 projected to the RN, 39 to the PN, and 155 neurons had no identifiable projection to these nuclei. No differences were observed in the recording depth among CR, CP, and non-identified neurons (CR = 785.02 ± 32.1 μm, CP = 812.8 ± 33.9 μm, NI = 778.2 ± 19 μm; F = 0.373, p = 0.689, One Ay Analysis of Variance). However, CR neurons had a higher conduction time (5.09 ± 0.3 ms) as compared to CP neurons (2.54 ± 0.1 ms) (U = 1553.0, p<0.001, Mann-Whitney U statistic) (Figure 3E). Moreover, the spontaneous firing rate of CR neurons was significantly higher than that of CP neurons (3.02 ± 0.41 Hz, 1.44 ± 0.17 Hz, respectively; U = 585.00, p = 0.002 Mann-Whitney rank sum test) and the action potential duration of CP neurons was significantly lower than that of CR neurons (peak-to-valley duration; CR = 0.85 ± 0.01 ms; CP neurons = 0.72 ± 0.01 ms; U = 409.500, p =<0.001, Mann-Whitney U test)

**Figure 3.**
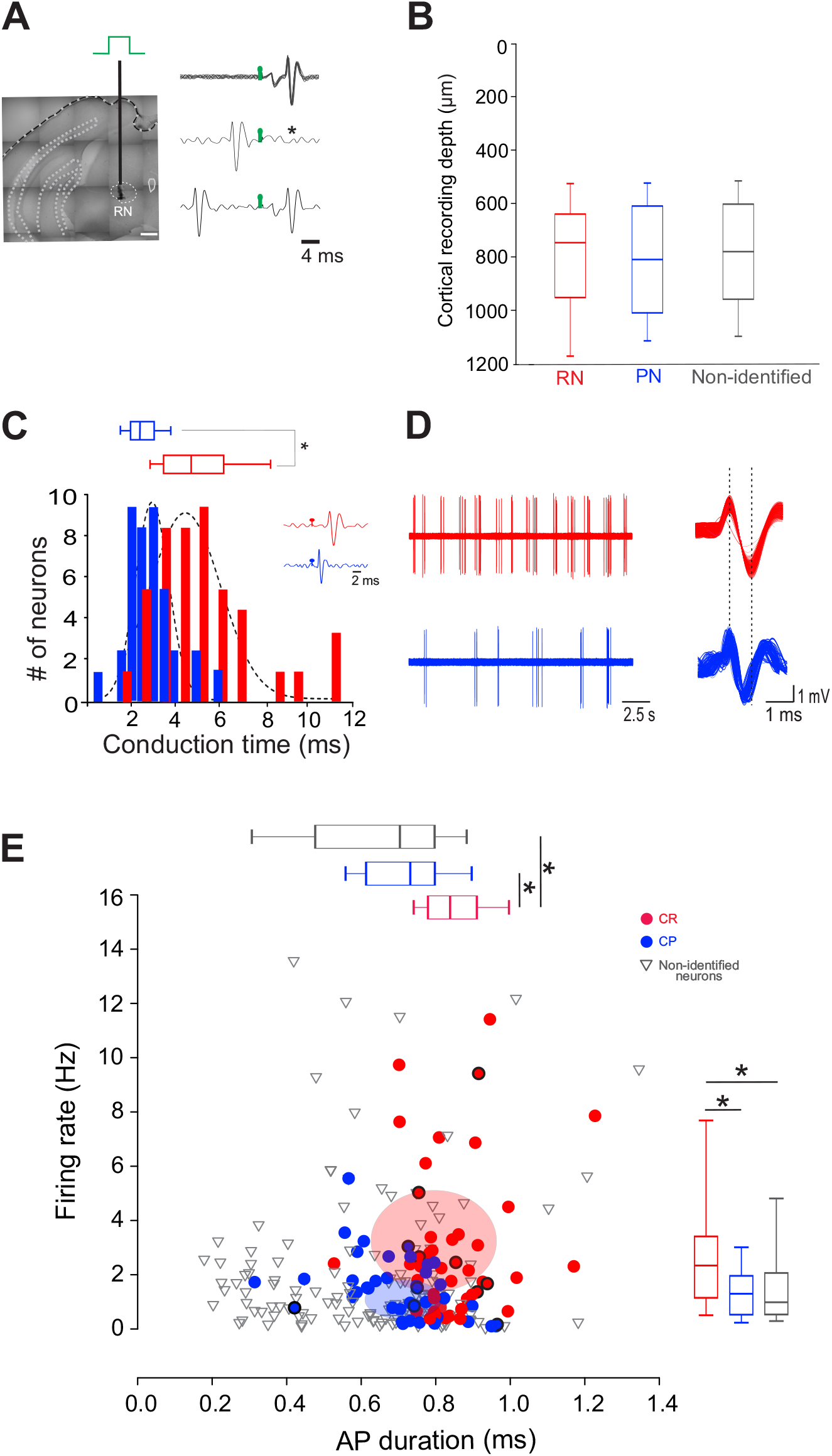
CR and CP PTNs exhibit distinct electrophysiological characteristics *in vivo*. **A**, PTNs were identified by collision tests. A stimulating electrode was placed in the RN or PN to stimulate the CR or CP terminals. The representative traces show superimposed electrophysiological antidromic responses evoked by the ipsilateral RN (upper trace) showing that action potentials appear with a fixed latency (stimulation artefact in green). The middle trace shows the collision test when an orthodromic action potential occurs below the antidromic latency. The asterisk indicates the point at which the antidromic response would have occurred. The third trace shows that when the orthodromic action potential occurs above the antidromic latency, the collision of antidromic response does not occur. **B**, Comparison between the recording depths obtained for PTNs projecting to RN, PN and non-identified neurons. **C**, Distribution of antidromic conduction time of CR (red) and CP (blue) neurons showing that neurons projecting to the RN conduct significantly faster (*p < 0.05, Mann-Whitney). Two exemplary traces shown in the inset. **D**, Representative electrophysiological recordings of spontaneous activity of CR (left upper trace) and CP (left lower trace). Aligned action potentials of the same neurons are also shown (right traces). **E**, Relationship between action potential (AP) duration (peak to valley) and spontaneous firing rate of all recorded neurons (CR n=45; CP n=42; non-identified n=154). Box plots show the distribution of AP durations (up) and firing rate (right). Ellipsoids in the graph represent the median (center) and standard errors (axis) of the data. *p < 0.05, One-way ANOVA, Dunn’s post hoc test and Shapiro-Wilk test.

### Axonal cortico-rubral and cortico-pontine projections display different conduction velocities

To analyze whether differences in antidromic conduction time are explained by differences in conduction velocity or merely a reflection of axonal length, we measured the length projections between M1 and RN or PN by means of virtual tractography (see Methods). The length of the axonal projections between M1 and either the ipsilateral RN or PN were estimated as the average length of ipsilateral virtual tracts reconstructed from 30 axonal tracing experiments from the Allen Mouse Brain Connectivity Atlas (Oh et al., 2014). Descending axons travel together by internal capsule reaching the cerebral peduncle. As reported previously in the rat (Brown et al., 1977), CR axons leave the cerebral peduncle and penetrate the substantia nigra and medial lemniscus to enter the RN (CR long). However, some fibers exit the cerebral peduncle in a rostral manner and penetrate the thalamus to reach the RN (CR short) (Figure 4). The distance traveled by CP axons (7452.03 ± 721.5 μm, n= 2305) is higher than that of CR bundles (CR long: 6754.99 ± 535.1 μm, n=61; CR short: 5865.43 ± 672.2 μm, n=44). With the measured mean pathway lengths of the tracts and the individual antidromic reaction times (Figure 3C), we computed the conduction velocities, which revealed that CP axons conduct significantly faster (3.64 ± 0.43 m/s) than CR axons (1.52 ± 0.11 m/s), without a difference between both pathways of CR neurons (H=68.8, p < 0.001, Kruskal-Wallis One Way ANOVA; Figure 4).

**Figure 4.**
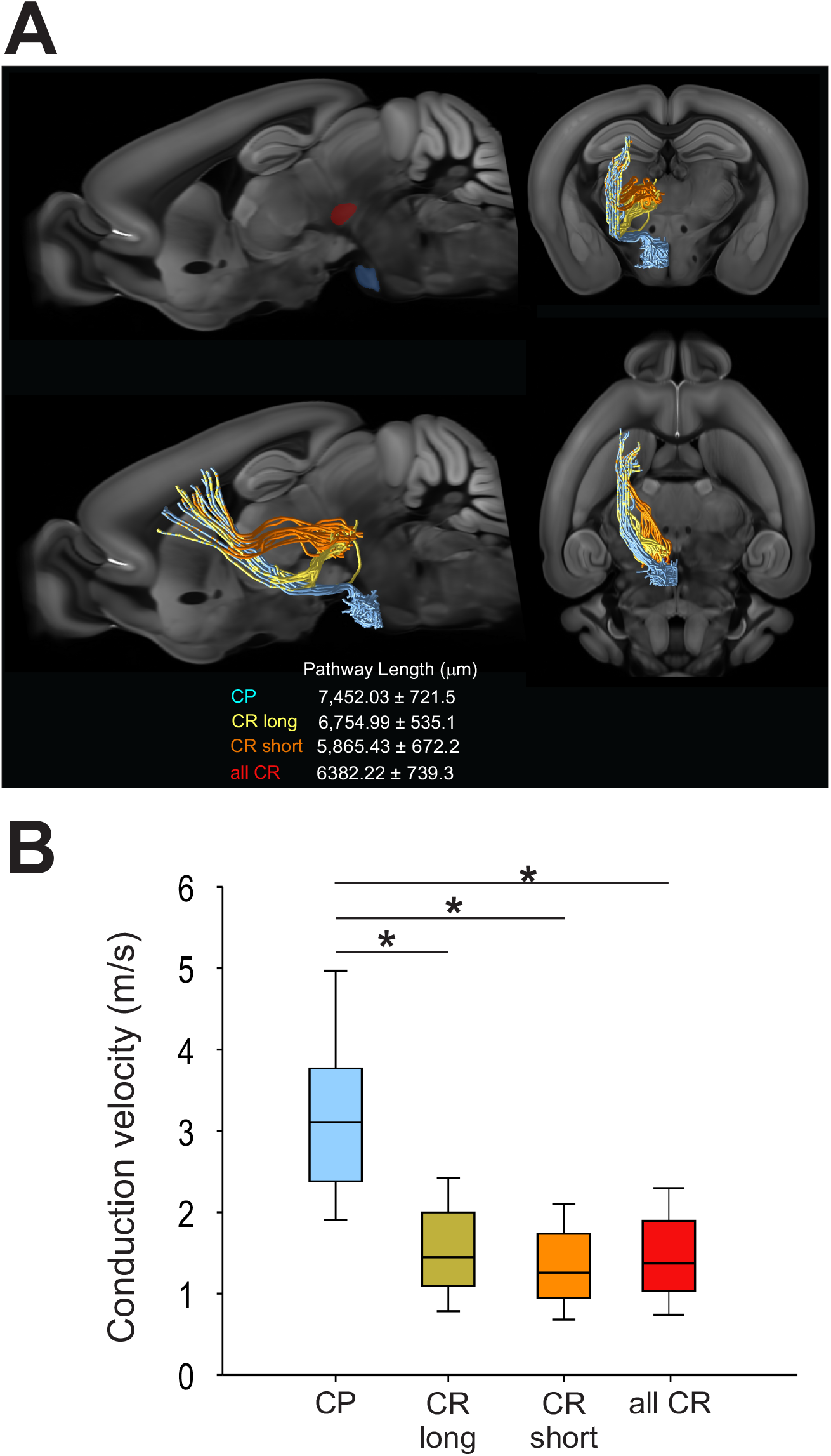
CR and CP neurons conduct at different velocities. **A**, Visual tractography reconstructed from 30 axonal tracing experiments from the Allen Mouse Brain Connectivity Atlas, showing the target ROIs and the pathway length from CP and CR axons traveling through the thalamus (orange CR short) or substantia nigra (yellow CR long) in the different anatomical planes. **B**, Comparison between the conduction velocities computed from conduction times (Figure 2C) and mean pathways lengths. Boxplots represents the median and 10-90 interquartile range. * p < 0.01 Kruskal-Wallis, One Way ANOVA, Dunn’s post hoc test.

### Morphology of cortico-rubral and cortico-pontine neurons

Some of the *in vivo* recorded neurons were filled with biocytin by means of juxtacellular recordings and microiontophoresis, allowing us to reconstruct the dendrite morphologies of 11 labeled PTNs: 6 CR and 5 CP (Figure 5A, E). All reconstructed neurons had the same coordinates within M1 (0.98 mm posterior and 1 mm lateral relative to bregma). The tracing results were aligned to the pial surface, which allowed us to determine the 3D soma, dendrite locations and dendritic spines with 50 μm precision. The results show that the basal dendrites of CR neurons are more extensive and more variable as those of CP neurons. These differences were quantified by comparing the distribution of the dendritic density (Figure 5B, F) and number of spines along the cortical depth (Figure 5D, H), which are different in CR and CP neurons (Dendrite density h = 1, p = <0.0049; Dendritic spine number h = 1, p= <0.001; Kolmogorov-Smirnov) Moreover, we quantified these differences by means of two indices to analyze the similarity (see Methods) between dendrite and spine density distributions of each individual neuron and the two dendrite or spine averaged distributions across PTNs with the same target (Figure 5J). The similarity analysis revealed that both dendrite and spine density distributions along cortical depth were more similar when PTNs had the same target (dendritic length similarity index: CP = -0.37 ± 0.07 μm vs CR = 0.24 ± 0.06, t = 6.168, p=<0.001; dendritic spine similarity index: CP = -0.18 ± 0.11 vs CR = 0.14 ± 0.07, t = 2.387, p<0.041 Student t-test).

**Figure 5.**
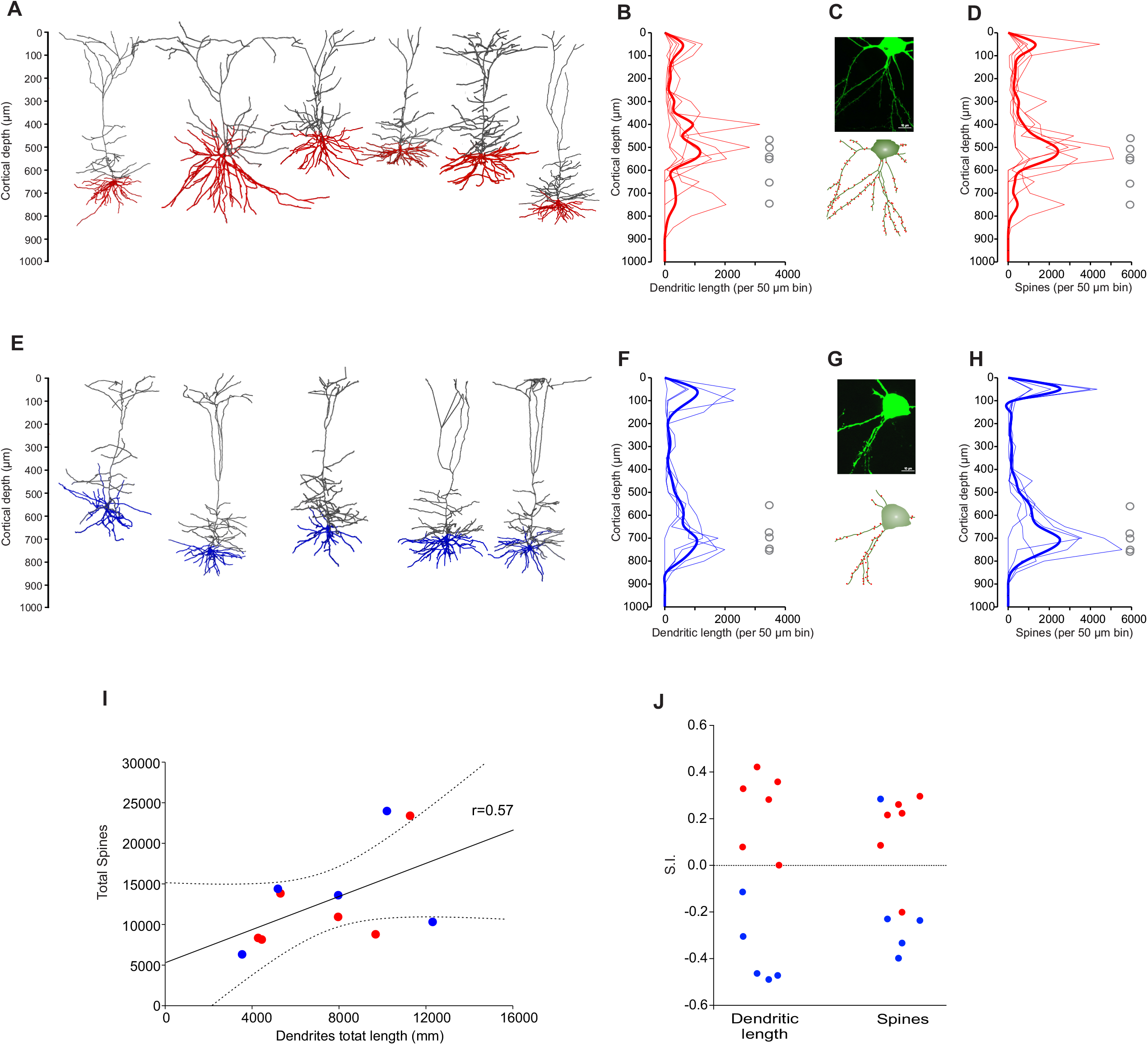
Dendrite morphology of PTNs reflects subcortical targets. **A**, Three-dimensional reconstructions of labeled CR neurons identified *in vivo* (n=6). Basal dendrites are indicated in red. **B**, Distribution of dendrites along the vertical cortex axis. Each thin line corresponds to one neuron, and the average with the standard error is indicated by a thick line and shadow. The circles represent soma depth of each neuron. **C**, Confocal microphotography of an exemplary CR neuron (upper) and its reconstruction (lower) showing the soma and some dendritic spines (red). **D**, Distribution of dendritic spines along the vertical axis of the cortex. Each thin line corresponds to one neuron, and the average with the standard error is indicated by a thick line and shadow. **E-H**, The same as A-D, but for CP neurons. **I**, Dendritic path length per PTN is significantly correlated to the respective number of dendritic spines. (CR red circles, CP blue circles). Lines represent the linear regression and the upper and lower 99% confidence intervals. **J**, Similarity indices (S.I.) between distributions along the cortex of dendrite density length and number of spines of each PTN. Notice that most of the similar neurons are of the same projection type.

For each PTN, we extracted a set of 9 morphological parameters (Table 2); no statistical differences between CR and CP neurons were found in any of the parameters. Additionally, the soma depth distributions of the CR- and CP-reconstructed neurons were similar to the overall distribution of the neurons labeled with retrograde tracers in M1 (range: 400-1100 μm, peak: 850 μm, Figure 2). Soma depth locations were not significantly different between reconstructed CR (n= 6; 600 ± 36.515 μm) and CP (n=5; 700 ± 27.386 μm) neurons (t = - 2.113 p = 0.064, Student t-test).

**Table 2.** Total characteristics of the projection neurons to red nucleus and pons.

| Dendritic Features | Corticorubral neurons (n=6) | Corticopontine neurons (n=5) | P value |
| --- | --- | --- | --- |
| **Cell Structure (mean±SD)** | **Total dendrites** |  |  |
| Total dendrites Length (μm) | 7157.16 ± 1197.28 | 7839.90 ± 1598.56 | 0.735 |
| Total nodes | 876.83 ± 80.85 | 1083.6 ± 162.09 | 0.298 |
| Intermediate nodes | 643.83 ± 66.12 | 846.4 ± 142.21 | 0.792 |
| Branching nodes | 111.16 ± 8.57 | 113.4 ± 12.25 | 0.891 |
| Terminal nodes | 121.83 ± 9.54 | 123.8 ± 13.62 | 0.914 |
| Segments number | 796.16 ± 46.05 | 1084 ± 162.17 | 0.125 |
| BoundingBox (mm<sup>3</sup>) | 0.05 ± 0.01 | 0.07 ± 0.01 | 0.102 |
| Spine number | 12229.83 ± 2397.03 | 13712.00 ± 2931.97 | 0.702 |
| Soma depth (μm) | 574.8 ± 43.7 | 679.9 ± 35.1 | 0.103 |
|  | **Apical dendrites** |  |  |
| Total dendrites Length (μm) | 5218.71 ± 1202.98 | 5610.70 ± 1272.61 | 0.828 |
| Total nodes | 437.66 ± 57.59 | 638 ± 112.5 | 0.160 |
| Intermediate nodes | 322.66 ± 43.97 | 499.4 ± 92.1 | 0.128 |
| Branching nodes | 46.83 ± 9.80 | 64.8 ± 11.71 | 0.303 |
| Terminal nodes | 61.5 ± 7.53 | 73.8 ± 11.87 | 0.427 |
| Segments number | 388.33 ± 37.60 | 636.6 ± 13.77 | 0.071 |
| Spine number | 8506.33 ± 1910.00 | 9126.40 ± 1907.82 | 0.662 |
|  | **Basal dendrites** |  |  |
| Total dendrites Length (μm) | 1938.45 ± 344.40 | 2229.20 ± 458.78 | 0.618 |
| Total nodes | 439.16 ± 33.84 | 445.6 ± 58.16 | 0.929 |
| Intermediate nodes | 321.16 ± 25.91 | 347 ± 57.04 | 0.931 |
| Branching nodes | 58.33 ± 7.29 | 48.6 ± 1.67 | 0.298 |
| Terminal nodes | 59.66 ± 6.39 | 50 ± 2.07 | 0.250 |
| Segments number | 412.83 ± 27.59 | 447.4 ± 56.79 | 0.792 |
| Spine number | 3723.50 ± 819.10 | 4585.60 ± 1287.18 | 0.573 |
| **Ongoing function (mean±SD)** |  |  |  |
| Firing rate (Hz) | 3.59 ± 1.21 | 1.26 ± 0.45 | 0.185 |
| Action potential duration (s) | $0.87 \pm 0.04$ | $0.72 \pm 0.08$ | 0.125 |
| Recording depth ($\mu\text{m}$) | $697.33 \pm 76.65$ | $841.2 \pm 43.87$ | 0.082 |

### *Cortico-rubral and cortico-pontine neurons* are necessary for proper movement execution

To understand the role of CR- and CP-descending commands on forelimb movements PTNs were optogenetically inhibited using the retrograde virus AAVrg-hsyn-Jaws-KGC-GFP-ER2 injected into the RN or PN of animals trained to press a lever in response to light (Figure 6A). An optical fiber was then implanted in ipsilateral M1; so that, CR or CP neurons expressing Jaws-ER2 halorhodopsin were inhibited with orange light (Chuong et al., 2014) during the task (see methods). No differences were found between the number of CP (356.1±26.05 neurons/ mm^2^) and CR (359.9±62.7 neurons/ mm^2^) expressing Jaws-ER2 halorhodopsin (t=0.0989; p=0.923). Photoinhibition of cortical neurons was performed during different movement phases, either before movement initiation or during movement execution (Figure 6B). The results show that the proportion of correct responses decreased significantly when CR or CP M1 neurons were inhibited before or during movements (CR control = 0.6 ± 0.02, before movement = 0.36 ± 0.41%, after movement = 0.33 ± 0.42 %, F = 16.45 p = 0.004 between groups, p = 0.005 between control vs 1; CP: control = 0.68 ± 0.01%, before movement = 0.49 ± 0.08 %, after movement = 0.300 ± 0.05 %; F= 10.703 p = 0.010 between groups, p= 0.001 between control vs 1). These results indicate that both cortico-mesencephalic projections participate in the execution of forelimb movements.

**Figure 6.**
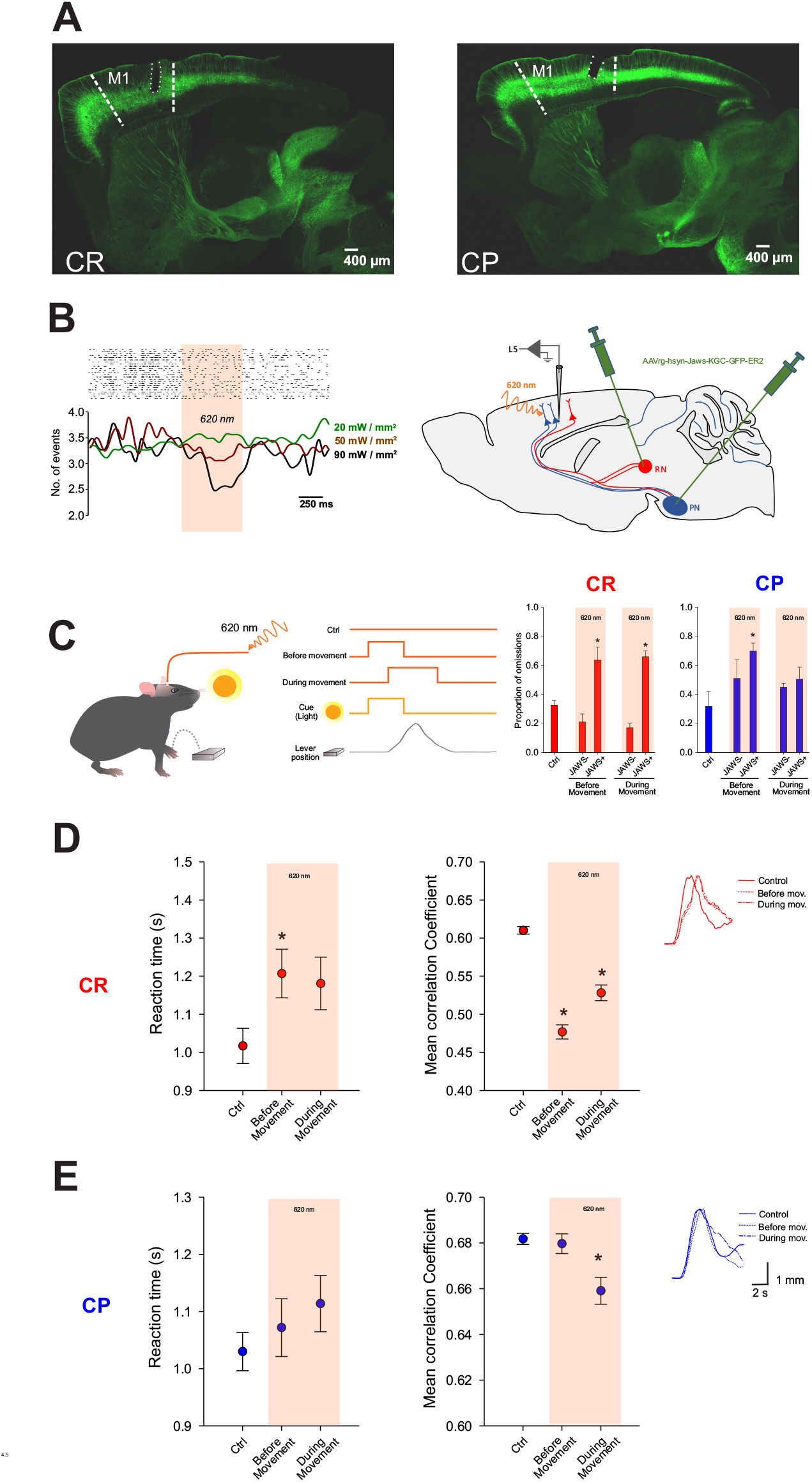
Specific inhibition of CR and CP PTNs affects movement onset and performance. **A**, Coronal microphotographs showing the expression of the Jaws retrograde virus (AAVrg-hsyn-Jaws-KGC-GFP-ER2) in CP (left) and CR (right) neurons, as well as the optical fiber location in M1. **B**, Raster display and firing spontaneous activity rate of an exemplary cortical neuron recorded during different light power intensity photoinhibition in an animal injected in the PN with Jaws retrograde virus. C, In trained animals, photoinhibition during the behavioral task was randomly applied in two different trials: 1) Before movement: ON starting with the light cue - OFF when animals press the lever (2 s in incorrect trials); 2) During movement: ON 800 ms after light cue starts with a duration of 2 s. Comparison between the omissions computed for control trials (Ctrl: without photoinhibition in animals expressing JAWS halorhodopsin; JAWS-: animals without the actuator) and photoinhibition trials (JAWS+: animals expressing JAWS halorhodopsin) of CR and CP neurons (n=3 mice per group, during 3 experimental sessions). D, Reaction times of correct lever pressings (middle) and correlation coefficients between individual lever pressing trajectories computed for control trials, and photoinhibition trials of CR neurons (n=3 mice, during 3 experimental sessions). The insert shows averaged paw trajectories performed during lever pressings in different trials. **E**, Same as D but photoinhibition was performed in CP neurons. * p < 0.05 Kruskal-Wallis, One Way ANOVA, Dunn’s post hoc test vs control group.

To gain insight into how movements are modulated by CR and CP neurons we compared reaction times and movement trajectories in control and photoinhibition conditions (Figure 6C-D; correct trials were analyzed). We found that reaction times increased, and movement trajectories were more variable when inhibiting of either CR or CP neurons. Interestingly, the inhibition of CR neurons before movement initiation had much stronger effect in movement variability as compared to CP inhibition. These results suggest that both classes of neurons might play different roles in movement onset and movement execution.

### Viral tracing of CR and CP neurons

To further investigate the collateralization of CR and CP neurons we used MouseLight (http://mouselight.janelia.org/) to evaluate the connections between primary motor cortex, RN and PN. From the 1,200 neurons traced and available in this resource, we selected those with (i) their axonal end-points located at the PN or RN, and (ii) whose soma was located in the primary motor cortex. These selection criteria yielded 22 neurons. Seven of those neurons (neurons ID’s AA1543, AA1043, AA0587, AA0584, AA0169, AA0134, AA0133) had terminal end points at both RN and PN. All neurons identified as reaching PN also had axonal end points at RN; conversely, no neurons were identified having terminal end points in RN but not in PN. (Selection criteria: axonal end points in either RN or PN, soma in Primary motor cortex; resulting neurons: AA1543, AA1542, AA1540, AA1539, AA1537, AA1050, AA1043, AA0927, AA0926, AA0923, AA0617, AA0587, AA0584, AA0261, AA0169, AA0135, AA0134, AA0133, AA0132, AA0131, AA0114).

Additionally, we analyzed the neuron terminations of CP neurons in the RN and vice versa, the terminations of CR neurons into the PN in our experiments in which control retrograde virus used for optogenetics inhibition (AAVrg-hSyn-EGFP) was injected in one of the targets (red or pons nuclei). In these experiments, GFP protein is expressed in all the projections of the infected neurons. We observed that in both, RN and PN, there are scarce terminals of the neurons infected by the virus injected into the PN and RN, respectively (Figure 7). Viral tracing suggests that CP and CR neurons are not completely independent and share some inputs.

**Figure 7.**
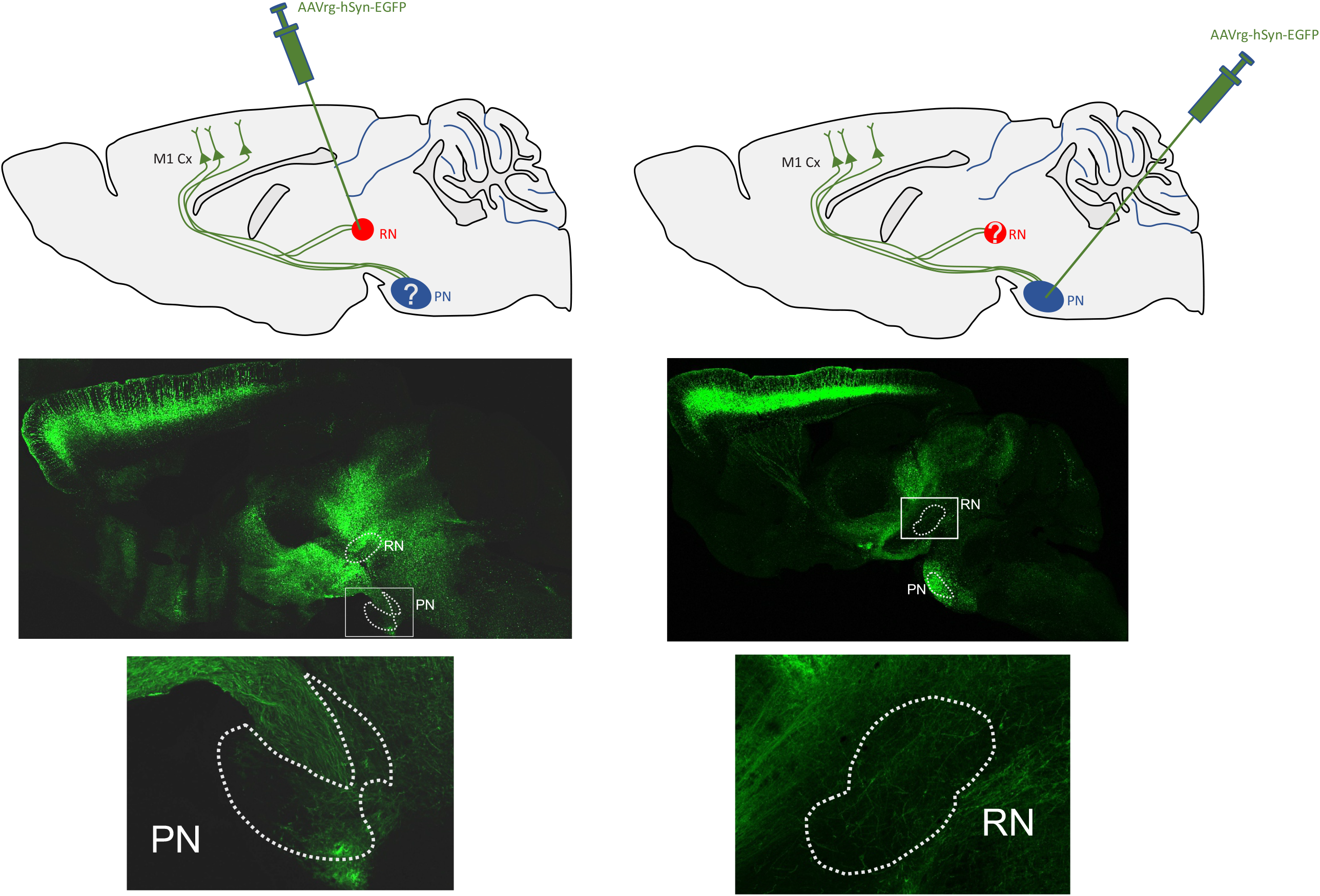
collateralization of CR and CP neurons. Retrograde virus (AAVrg-hSyn-EGFP) was injected in RN (left) and PN (right). Microphotographs of exemplary experiments showing neuron terminations of CR neurons (left) in the PN and CP neurons in the RN (right).

## DISCUSSION

We provided functional, anatomical, and morphological evidence indicating that two distinct populations of PTNs differentially contribute to motor execution. Previous studies have described the relationships between structure and function in Layer 5 PTNs projecting to different targets (Groh et al., 2010; Rojas-Piloni et al., 2017; Olivares-Moreno et al., 2019). Our data revealed that both classes of PTNs are intermingled in sensorimotor cortices, including M1, and only a small proportion of neurons seem to contact the RN and PN simultaneously. This does not imply that PTN endings arrive only at a single subcortical area; for example, in the mouse barrel field it has been reported that around 40% of PTNs project to two targets, and 20% to three targets including the striatum, thalamus, hypothalamus, midbrain, pons and spinal cord (Guo et al., 2017). Moreover, Rojas-Piloni et al. (2017) reported that ∼80% of all PTNs in the barrel field project only to 4 targets (posterior medial division of the thalamus, tectum, pontine nucleus and subnucleus caudalis of the spinal trigeminal tract) and ∼20% of them project to two targets. Here, we confirm that the number of neurons projecting simultaneously to the RN and PN is considerably smaller (Akintunde and Buxton, 1992). However, it is to be expected that the majority of CR and CP PTNs innervate additional targets not explored in our study. Quantification of collateralization is always a crucial problem and usually, the double-labeled cells in multiple retrograde tracing experiments are underestimated (Varcelli et al., 2000). The viral tracing experiments show that a major proportion of CP and CR neurons project to red and pontine nuclei, simultaneously (Economo et al., 2016). The difference observed between retrograde tracers and viral anterograde tracing reconstructions could be explained by the different mechanisms by which fluorescent molecules are expressed into the cells. Retrograde fluorescent tracers are taken by endocytosis at the synapses and only transported retrogradely to the soma; however, neurons infected by the viral vectors start to express fluorescent proteins revealing the neuron morphology and anatomical projections. In this way, is tempting to propose that retrograde tracing is a method reflecting not only the anatomic projections, but also pathway function.

Layer 5 thick-tufted PTNs are a canonical class of cells present in all neocortical areas (Ramaswamy and Markram, 2015) but also constitute the most morphologically and functionally variable type of neuron (Oberlaender et al., 2011; Kim et al., 2015; Rojas-Piloni et al., 2017). Here we found that CR and CP PTNs exhibit particular electrophysiological characteristics; that is, CR neurons have a higher firing rate and extracellular spike durations than CP neurons. This result confirms the previous findings showing that spike duration increases with increasing firing rate (Bourque and Renaud, 1985; Stratton et al., 2012). Both classes of neurons display different ongoing firing rates yet are in the same range of the previously reported PTNs in rats (Oberlaender et al., 2011; Rojas-Piloni et al., 2017).

In addition, we found that the antidromic spiking latency (conduction time) of CP neurons is shorter in CR neurons, suggesting a higher conduction velocity for neurons projecting to the pons. Using virtual tractography we demonstrated different conduction velocities for CR and CP neurons (1.5 and 3.6 m/s, respectively: range 1.3-18.6). The mean conduction velocities computed here are slightly less than conduction velocities reported in mice for the pyramidal tract (8.89±1.81 m/s) (Tanaka et al., 2004), as well as for corticospinal tract (13.7 m/s) (Powers et al 2012). Nevertheless, the discrepancies in conduction velocity estimated from anatomical or electrophysiological data have been explained by a recording bias (Towe and Harding 1970; Kraskov et al., 2020) or by the precision to measure the distance traveled by the axons. By enriching our electrophysiological recordings with the extensive axonal tracing data set available through the Allen Brain Connectivity Atlas (Oh et al., 2014), we show a clear difference in conduction velocities between M1 neurons projecting to the RN and PN. The differences in conduction velocities might arise from different myelination properties of both classes of PTNs. In fact, a great variability in the longitudinal distribution of PTN myelin has been reported, suggesting the presence of a neuronal identity that could be used as a strategy to modulate long-distance communication in the neocortex (Tomassy et al., 2014). Further, CP conduction times display less variability and more homogeneous distribution between recorded neurons (Figure 2C), implying that the information arrives with a precise timing. This potentially plays a role in sensorimotor integration the main target of pontocerebellar projections are granule cells, which receive inputs also from the brainstem and spinal cord (Sillitoe et al., 2012; Huang et al., 2013), and have been considered as coincidence detectors (Wagner et al., 2019).

Both CR and CP neurons are widely distributed into the sensorimotor cortices. Surprisingly, here we found behavioral significant effects using photoinhibition of a small area in M1 Photostimulation of small areas has been observed previously (Magno et al., 2019; Mathis et al, 2017). Additionally, microstimulation experiments have established a direct link between sensorimotor activity to behavior suggesting that modulation of a small neuronal population can influence sensory decisions (Houweling and Bretch, 2008).

When we analyzed individual 3D morphologies of PTNs projecting to the RN and PN, no differences were found between global characteristics measured in both classes (Table 2). However, the distribution of dendrites and dendritic spines along the cortex is clearly different between CR and CP neurons, but similar between individuals of each class. Since layer-specific axon innervation is a feature of cortical organization, observed for local and long-range axons (Wimmer et al., 2010; Narayanan et al., 2015), the particularities in laminar distribution of target-related dendrites might suggest differences in the synaptic input that each PTN class receives. Moreover, these differences can produce distinct spiking patterns in both classes of PTNs. Due to the complex interaction between the dendritic and axonal initiation in different dendritic subdomains (apical vs basal) of the same cell, the firing patterns of PTNs can be modulated by synchronous synaptic inputs arriving at different lamina (Larkum et al., 1999).

### Functional implications

Even though the role of the RN and PN in movement has been extensively described individually (Tziridis et al., 2009; Wagner et al., 2019; Basile et al., 2021), less is known about the specific contributions of each projection class. The function of the magnocellular part of the RN (that originates the rubrospinal tract) in movement has been well studied in behaviors like scratching, locomotion, and learned or automated motor behaviors (Kennedy, 1990; Gruber and Gould, 2010). However, there is little information about the parvocellular area that receives inputs from the cortex and originates rubro-olivary tracts. Lesions of the parvocellular region create mobility problems in rats, as it was shown that they could not perform precise movements, such as supination and pronation of the paw. They also exhibited deficiencies in digit movements (e.g., arpeggio) and had difficulty coordinating catching, aiming, and reaching efficiently (Whishaw et al., 1990; Whishaw et al., 1998; Morris et al., 2011; Morris et al., 2015). In agreement, here we found that inhibition of CR neurons produces important alterations of trajectories in stereotyped learned movements.

On the other hand, the PN also routes cerebral cortex information to the cerebellum (Brodal and Bjaalie, 1992; Wagner et al., 2019). PN unitary neuronal responses during forelimb movements in monkeys (Tziridis et al., 2009) have been recorded, and PN lesions in humans produce motor deficits (Schmahmann et al., 2004)). A recent study tested the role of the PN in the execution of cortically dependent voluntary movements and indicated that it is essential for dexterous forelimb motor control (Guo et al., 2019).

We found that both CR and CP neurons increase their activity after a visual cue and movement initiation, but CP neurons activate earlier than CR neurons. It has been reported that the neurons of these nuclei also display responses to sensory stimulation; PN neurons respond to visual and auditory stimulation (Campolattaro et al., 2011), whereas RN neurons mainly respond to somatosensory stimulation (Nishioka and Nakahama, 1973) have also been implicated in visually guided movement responses (Smith, 1970). The specific contribution of CR and CP PTNs to forelimb movement performance remains to be determined. The fact that correlation between individual movement trajectories decreased during inhibition of CR and CP PTNs indicates that the information propagated by both descending projections is necessary for proper movement performance. Nevertheless, reaction time increased only during CR inhibition (Figure 6), which is in line with classical descriptions showing that RN neurons modulated its activity during the onset of movements (Jarratt and Hyland, 1999). Moreover, there are significant differences in response time and movement variability when red nucleus projecting PTNs are inhibited before movement. These differences are not present when pons projecting PTNs are inhibited before movement, suggesting that the two groups have different contributions during the behavior. Because we only analyzed movement performed in the remaining correct responses, the results suggest that movement initiation depends more on corticopontine, but movement performance on corticorubral. Together with the electrophysiological and photometry experiments, it is possible to reinforce the idea that both classes of neurons process and conduct different information for sensorimotor integration.

We demonstrate that the activity of both classes of neurons increases long before movement onset (∼500 ms). Yet the neural activity of pons-projecting cortical neurons shows a significant correlation with the future movement (i.e., the M1 CP neurons synchronize their activity with a fixed time lag before movement initiates). This preparatory activity is present in several cortical areas and has been associated with motor planning, which implies moving the trigger networks to an initial condition that is matched to specific movements (Svoboda and Li, 2018). It remains to be determined if the difference in calcium peak timing displayed by PTNs projecting to the RN and PN before movement is due to differences in the firing frequency of the neurons. The calcium signal in neurons is secondary to the generation of action potentials, and its peak depends on the firing frequency of the neurons. The faster peak of calcium signal in CP neurons compared to CR neurons, may reflect not only their faster response but a higher frequency of their response compared to CR. Nevertheless, the functional and morphological differences in PTNs projecting to the RN and PN suggests that the sensorimotor cortex generates different downstream coding with separable contributions to volitive movements.

## ACKNOWLEDGEMENTS

We thank Dr. Marcel Oberlaender for his critical feedback on this manuscript; Jessica Gonzalez Norris for proofreading the manuscript; and Cutberto Dorado, Nydia Hernandez, Ericka de los Rios, Alejandra Castilla, Adriana Gonzalez Gallardo, Martín Garcia Servin and Christian Josue Delgado Guzman for providing technical assistance. VL-V, PR-M, and MM are doctoral students from the Programa de Doctorado en Ciencias Biomédicas, Universidad Nacional Autónoma de México (UNAM) and received fellowships from CONACyT (660523, 934973, and 858704). This work was funded by grants from CONACYT Ciencia Básica A1-S-8686 (GR-P), UNAM-DGAPA-PAPIME PE205821 (RO-M), and UNAM-DGAPA PAPIIT IN201121 (GR-P).

## FIELD STATEMENT

Pyramidal tract neurons (PTNs) of the sensorimotor cortex are fundamental for execution of movements. Because they project to several subcortical structures, a fundamental question is whether PTNs are organized, generating different and specific output commands for sensorimotor integration. This idea has long been suggested but it is largely unknown. Here, we have shown evidence supporting that PTNs segregates into different functional subsystems integrating and conducting in a coordinated manner distinct information related to sensorimotor information. This means that functional compartmentalization and hierarchical organization exist between PTNs projecting to distinct subcortical zones.

## REFERENCES

Akintunde A, Buxton DF (1992). Origins and collateralization of corticospinal, corticopontine, corticorubral and corticostriatal tracts: a multiple retrograde fluorescent tracing study. Brain Res 586:208–218. doi:10.1016/0006-8993(92)91629-S.

Angaut P, Bowsher D (1965). Cerebello-rubral Connexions in the Cat. Nature 208:1002–1003. doi: 10.1038/2081002a0

Basile GA, Quartu M, Bertino S, Serra MP, Boi M, Bramanti A, Anastasi GP, Milardi D, Cacciola A (2021) Red nucleus structure and function: from anatomy to clinical neurosciences. Brain structure & function 226:69–91. doi: 10.1007/s00429-020-02171-x

Bourque CW, Renaud LP (1985) Activity dependence of action potential duration in rat supraoptic neurosecretory neurones recorded in vitro. J Physiol 363:429–439.doi: 10.1113/jphysiol.1985.sp015720

Brodal P (2014) Pontine Nuclei. In: Encyclopedia of the Neurological Sciences (Second Edition) (Aminoff MJ, Daroff RB, eds), pp 938–940. Oxford: Academic Press.

Brodal P, Bjaalie JG (1992) Organization of the pontine nuclei. Neuroscience Research 13:83–118. doi: 10.1016/0168-0102(92)90092-Q

Brown JT, Chan-Palay V, Palay SL (1977) A study of afferent input to the inferior olivary complex in the rat by retrograde axonal transport of horseradish peroxidase. Journal of Comparative Neurology 176:1–22. doi: 10.1002/cne.901760102

Campolattaro MM, Kashef A, Lee I, Freeman JH (2011) Neuronal Correlates of Cross-Modal Transfer in the Cerebellum and Pontine Nuclei. The Journal of Neuroscience 31:4051–4062. doi: 10.1523/JNEUROSCI.4142-10.2011

Catman-Berrevoets CE, Kuypers HGJM, Lemon RN (1979) Cells of origin of the frontal projections to magnocellular and parvocellular red nucleus and superior colliculus in cynomolgus monkey. An HRP study. Neuroscience Letters 12:41–46. doi: 10.1016/0304-3940(79)91477-0

Courville J (1966) Somatotopical organization of the projection from the nucleus interpositus anterior of the cerebellum to the red nucleus. An experimental study in the cat with silver impregnation methods. Experimental Brain Research 2:191–215. doi: 10.1007/BF00236713

Chuong AS et al. (2014) Noninvasive optical inhibition with a red-shifted microbial rhodopsin. Nature neuroscience 17:1123–1129. doi: 10.1038/nn.3752

Daniel H, Angaut P, Batini C, Billard JM (1988) Topographic organization of the interpositorubral connections in the rat. A WGA-HRP study. Behavioural Brain Research 28:69–70. doi: 10.1016/0166-4328(88)90078-2

Economo MN, Clack NG, Lavis LD, Gerfen CR, Svoboda K, Myers EW, Chandrashekar J (2016) A platform for brain-wide imaging and reconstruction of individual neurons eLife 5:e10566. doi: 10.7554/eLife.10566

Economo MN, Viswanathan S, Tasic B, Bas E, Winnubst J, Menon V, Graybuck LT, Nguyen TN, Smith KA, Yao Z, Wang L, Gerfen CR, Chandrashekar J, Zeng H, Looger LL, Svoboda K (2018) Distinct descending motor cortex pathways and their roles in movement. Nature 563:79–84. doi: 10.1038/s41586-018-0642-9

Edwards SB (1972) The ascending and descending projections of the red nucleus in the cat: An experimental study using an autoradiographic tracing method. Brain Research 48:45–63. doi: 10.1016/0006-8993(72)90170-9

Groh A, Meyer HS, Schmidt EF, Heintz N, Sakmann B, Krieger P (2010) Cell-type specific properties of pyramidal neurons in neocortex underlying a layout that is modifiable depending on the cortical area. Cereb Cortex 20:826–836. doi: 10.1093/cercor/bhp152

Gruber P, Gould DJ (2010) The Red Nucleus: Past, Present, and Future. Neuroanatomy 9:1–3.

Guo C, Peng J, Zhang Y, Li A, Li Y, Yuan J, Xu X, Ren M, Gong H, Chen S (2017) Single-axon level morphological analysis of corticofugal projection neurons in mouse barrel field. Scientific Reports 7:2846. doi: 10.1038/s41598-017-03000-8

Guo J-Z, Sauerbrei B, Cohen JD, Mischiati M, Graves A, Pisanello F, Branson K, Hantman AW (2019) The Pontine Nuclei are an Integrative Cortico-Cerebellar Link Critical for Dexterity. bioRxiv:637447. doi: 10.1101/637447

Houweling AR, Brecht M (2008) Behavioural report of single neuron stimulation in somatosensory cortex. Nature. 451(7174):65–68. doi: 10.1038/nature6447

Huang CC, Sugino K, Shima Y, Guo C, Bai S, Mensh BD, Nelson SB, Hantman AW (2013) Convergence of pontine and proprioceptive streams onto multimodal cerebellar granule cells. eLife 2013. doi: 10.7554/eLife.00400

Humphrey DR, Rietz RR (1976) Cells of origin of corticorubral projections from the arm area of primate motor cortex and their synaptic actions in the red nucleus. Brain Research 110:162–169. doi: 10.1016/0006-8993(76)90217-1

Jansen J, Jansen Jr J (1955) On the efferent fibers of the cerebellar nuclei in the cat. Journal of Comparative Neurology 102:607–632. doi: 10.1002/cne.901540202

Janke AL, Ullmann JF. (2015) Robust methods to create ex vivo minimum deformation atlases for brain mapping. Methods. 2015 Feb;73:18–26. doi: 10.1016/j.ymeth.2015.01.005

Jarratt H, Hyland B (1999) Neuronal activity in rat red nucleus during forelimb reach-to-grasp movements. Neuroscience 88:629–642. doi: 10.1016/S0306-4522(98)00227-9

Jiang S, Guan Y, Chen S, Jia X, Ni H, Zhang Y, Han Y, Peng X, Zhou C, Li A, Luo Q, Gong H (2020) Anatomically revealed morphological patterns of pyramidal neurons in layer 5 of the motor cortex. Scientific Reports 10:7916. doi: 10.1038/s41598-020-64665-2

Kennedy PR (1990) Corticospinal, rubrospinal and rubro-olivary projections: a unifying hypothesis. Trends in Neurosciences 13:474–479. doi: 10.1016/0166-2236(90)90079-p

Kim EJ, Juavinett AL, Kyubwa EM, Jacobs MW, Callaway EM (2015) Three Types of Cortical Layer 5 Neurons That Differ in Brain-wide Connectivity and Function. Neuron 88:1253–1267. doi: 10.1016/j.neuron.2015.11.002

Kita T, Kita H (2012) The Subthalamic Nucleus Is One of Multiple Innervation Sites for Long-Range Corticofugal Axons: As Single-Axon Tracing Study in the Rat. The Journal of Neuroscience 32:5990–5999. doi: 10.1523/JNEUROSCI.5717-11.2012

Kraskov A, Soteropoulos DS, Glover IS, Lemon RN, Baker SN (2020) Slowly-Conducting Pyramidal Tract Neurons in Macaque and Rat. Cereb Cortex. 30(5):3403–3418. doi: 10.1093/cercor/bhz318

Larkum ME, Zhu JJ, Sakmann B (1999) A new cellular mechanism for coupling inputs arriving at different cortical layers. Nature 398:338–341. doi: 10.1038/18686

Lemon RN (2016) Cortical projections to the red nucleus and the brain stem in the rhesus monkey. Brain Res 1645:28–30. doi: 10.1016/j.brainres.2016.01.006

Liang H, Paxinos G, Watson C (2012) The red nucleus and the rubrospinal projection in the mouse. Brain Structure and Function 217:221–232. doi: 10.1007/s00429-011-0348-3

Magno LAV, Tenza-Ferrer H, Collodetti M, Aguiar MFG, Rodrigues APC, da Silva RS, Silva JDP, Nicolau NF, Rosa DVF, Birbrair A, Miranda DM, Romano-Silva MA (2019) Optogenetic Stimulation of the M2 Cortex Reverts Motor Dysfunction in a Mouse Model of Parkinson’s Disease. J Neurosci. 39(17):3234–3248. doi: 10.1523/JNEUROSCI.2277-18.2019

Massion J (1967) The mammalian red nucleus. Physiological Reviews 47:383–436. doi: 10.1152/physrev.1967.47.3.383

Mathis MW, Mathis A, Uchida N (2017) Somatosensory Cortex Plays an Essential Role in Forelimb Motor Adaptation in Mice. Neuron. 93(6):1493–1503.e6. doi: 10.1016/j.neuron.2017.02.049

Morris R, Tosolini AP, Goldstein JD, Whishaw IQ (2011) Impaired arpeggio movement in skilled reaching by rubrospinal tract lesions in the rat: a behavioral/anatomical fractionation. J Neurotrauma 28:2439–2451. doi: 10.1089/neu.2010.1708

Morris R, Vallester KK, Newton SS, Kearsley AP, Whishaw IQ (2015) The differential contributions of the parvocellular and the magnocellular subdivisions of the red nucleus to skilled reaching in the rat. Neuroscience 295:48–57. doi: 10.1016/j.neuroscience.2015.03.027

Narayanan, R.T., Mohan, H., Broersen, R., de Haan, R., Pieneman, A.W., de Kock, C.P., 2014. Juxtasomal biocytin labeling to study the structure-function relationship of individual cortical neurons. J. Vis. Exp.(84), e51359. doi: 10.3791/51359.

Narayanan RT, Egger R, Johnson AS, Mansvelder HD, Sakmann B, de Kock CPJ, Oberlaender M (2015) Beyond Columnar Organization: Cell Type-and Target Layer-Specific Principles of Horizontal Axon Projection Patterns in Rat Vibrissal Cortex. Cerebral cortex (New York, NY : 1991) 25:4450–4468. doi: 10.1093/cercor/bhv053

Nishioka S, Nakahama H (1973) Peripheral somatic activation of neurons in the cat red nucleus. Journal of Neurophysiology 36:296–307. doi: 10.1152/jn.1973.36.2.296

Oberlaender M, Bruno RM, Sakmann B, Broser PJ (2007) Transmitted light brightfield mosaic microscopy for three-dimensional tracing of single neuron morphology. Journal of biomedical optics 12:064029. doi: 10.1117/1.2815693

Oberlaender M, Boudewijns ZSRM, Kleele T, Mansvelder HD, Sakmann B, de Kock CPJ (2011) Three-dimensional axon morphologies of individual layer 5 neurons indicate cell type-specific intracortical pathways for whisker motion and touch. Proceedings of the National Academy of Sciences 108:4188–4193. doi: 10.1073/pnas.1100647108

Oh SW et al. (2014) A mesoscale connectome of the mouse brain. Nature 508:207–214. doi: 10.1038/nature13186

Olivares-Moreno R, Rodriguez-Moreno P, Lopez-Virgen V, Macías M, Altamira-Camacho M, Rojas-Piloni G (2021) Corticospinal vs Rubrospinal Revisited: An Evolutionary Perspective for Sensorimotor Integration. Frontiers in Neuroscience 15. doi: 10.3389/fnins.2021.686481

Olivares-Moreno R, Moreno-Lopez Y, Concha L, Martínez-Lorenzana G, Condés-Lara M, Cordero-Erausquin M, Rojas-Piloni G (2017) The rat corticospinal system is functionally and anatomically segregated. 222:3945–3958. doi: 10.1007/s00429-017-1447-6

Olivares-Moreno R, López-Hidalgo M, Altamirano-Espinoza A, González-Gallardo A, Antaramian A, Lopez-Virgen V, Rojas-Piloni G (2019) Mouse corticospinal system comprises different functional neuronal ensembles depending on their hodology. BMC Neuroscience 20:50. doi: 10.1186/s12868-019-0533-5

Oswald M, Tantirigama M, Sonntag I, Hughes S, Empson R (2013) Diversity of layer 5 projection neurons in the mouse motor cortex. Frontiers in Cellular Neuroscience 7. doi: 10.3389/fncel.2013.00174

Pinault D (1996) A novel single-cell staining procedure performed in vivo under electrophysiological control: morpho-functional features of juxtacellularly labeled thalamic cells and other central neurons with biocytin or Neurobiotin. J. Neurosci. Methods 65, 113–136. https://doi.org/10.1016/0165-0270(95)00144-1. doi: 10.1016/0165-0270(95)00144-1

Powers BE, Lasiene J, Plemel JR, Shupe L, Perlmutter SI, Tetzlaff W, Horner PJ. (2012) Axonal thinning and extensive remyelination without chronic demyelination in spinal injured rats. J Neurosci. 32(15):5120–5. doi: 10.1523/JNEUROSCI.0002-12.2012

Ramaswamy S, Markram H (2015) Anatomy and physiology of the thick-tufted layer 5 pyramidal neuron. Front Cell Neurosci 9:233. doi: 10.3389/fncel.2015.00233

Rojas-Piloni G, Guest JM, Egger R, Johnson AS, Sakmann B, Oberlaender M (2017) Relationships between structure, in vivo function and long-range axonal target of cortical pyramidal tract neurons. Nature Communications 8:870. doi: 10.1038/s41467-017-00971-0

Schmahmann JD, Ko R, MacMore J (2004) The human basis pontis: motor syndromes and topographic organization. Brain 127:1269–1291. doi: 10.1093/brain/awh138

Schwarz C, Thier P (1995) Modular organization of the pontine nuclei: dendritic fields of identified pontine projection neurons in the rat respect the borders of cortical afferent fields. The Journal of neuroscience : the official journal of the Society for Neuroscience 15:3475–3489. doi: 10.1523/JNEUROSCI.15-05-03475.1995

Sillitoe RV, Fu Y, Watson C (2012) Chapter 11 - Cerebellum. In: The Mouse Nervous System (Watson C, Paxinos G, Puelles L, eds), pp 360–397. San Diego: Academic Press.

Smith AM (1970) Deficits in conditioned movement and visual discrimination following rubral area lesions in the rat. Physiology & Behavior 5:893–896. doi: 10.1016/0031-9384(70)90178-2

Stratton P, Cheung A, Wiles J, Kiyatkin E, Sah P, Windels F (2012) Action potential waveform variability limits multi-unit separation in freely behaving rats. PLoS One 7:e38482. doi: 10.1371/journal.pone.0038482

Svoboda K, Li N (2018) Neural mechanisms of movement planning: motor cortex and beyond. Current opinion in neurobiology 49:33–41. doi: 10.1016/j.conb.2017.10.023

Tan, PN. (2009). Receiver Operating Characteristic. In: Liu, L., ÖZSU, M.T. (eds) Encyclopedia of Database Systems. Springer, Boston, MA. doi: 10.1007/978-0-387-39940-9_569

Tanaka H, Ono K, Shibasaki H, Isa T, Ikenaka K (2004) Conduction properties of identified neural pathways in the central nervous system of mice in vivo. Neurosci Res. 49(1):113–22. doi: 10.1016/j.neures.2004.02.001

Tomassy GS, Berger DR, Chen HH, Kasthuri N, Hayworth KJ, Vercelli A, Seung HS, Lichtman JW, Arlotta P (2014) Distinct profiles of myelin distribution along single axons of pyramidal neurons in the neocortex. Science 344:319–324. doi: 10.1126/science.1249766

Tournier JD, Smith R, Raffelt D, Tabbara R, Dhollander T, Pietsch M, Christiaens D, Jeurissen B, Yeh CH, Connelly A (2019) MRtrix3: A fast, flexible and open software framework for medical image processing and visualisation. NeuroImage 202:116137. doi: 10.1016/j.neuroimage.2019.116137

Towe AL, Harding GW (1970) Extracellular microelectrode sampling bias. Experimental Neurology 29:366–381. doi: 10.1016/0014-4886(70)90065-8

Tziridis K, Dicke PW, Thier P (2009) The Role of the Monkey Dorsal Pontine Nuclei in Goal-Directed Eye and Hand Movements. The Journal of Neuroscience 29:6154–6166. doi: 10.1523/JNEUROSCI.0581-09.2009

Ullmann JF, Janke AL, Reutens D, Watson C (2015) Development of MRI-based atlases of non-human brains. The Journal of comparative neurology 523:391–405. doi: 10.1002/cne.23678

Vercelli A, Repici M, Garbossa D, Grimaldi A (2000) Recent techniques for tracing pathways in the central nervous system of developing and adult mammals. rain Research Bulletin 51:11–28. doi: 10.1016/s0361-9230(99)00229-4

Wagner MJ, Kim TH, Kadmon J, Nguyen ND, Ganguli S, Schnitzer MJ, Luo L (2019) Shared Cortex-Cerebellum Dynamics in the Execution and Learning of a Motor Task. Cell 177:669–682.e624. doi: 10.1016/j.cell.2019.02.019

Walberg F, Nordby T (1981) A re-examination of the rubro-olivary tract in the cat, using horseradish peroxidase as a retrograde and an anterograde neuronal tracer. Neuroscience 6:2379–2391. doi: 10.1016/0306-4522(81)90024-5

Wang Q et al. (2020) The Allen Mouse Brain Common Coordinate Framework: A 3D Reference Atlas. Cell 181:936–953.e920. doi: 10.1016/j.cell.2020.04.007

Whishaw IQ, Tomie J-A, Ladowsky RL (1990) Red nucleus lesions do not affect limb preference or use, but exacerbate the effects of motor cortex lesions on grasping in the rat. Behavioural Brain Research 40:131–144. doi: 10.1016/0166-4328(90)90005-y

Whishaw IQ, Gorny B, Sarna J (1998) Paw and limb use in skilled and spontaneous reaching after pyramidal tract, red nucleus and combined lesions in the rat: behavioral and anatomical dissociations. Behav Brain Res 93:167–183. doi: 10.1016/s0166-4328(97)00152-6

Wimmer VC, Bruno RM, de Kock CPJ, Kuner T, Sakmann B (2010) Dimensions of a Projection Column and Architecture of VPM and POm Axons in Rat Vibrissal Cortex. Cerebral Cortex 20:2265–2276. doi: 10.1093/cercor/bhq068

Winnubst J et al. (2019) Reconstruction of 1,000 Projection Neurons Reveals New Cell Types and Organization of Long-Range Connectivity in the Mouse Brain. Cell 179:268–281.e213. doi: 10.1016/j.cell.2019.07.042

